# Voluntary muscle activation in people with Multiple Sclerosis is reduced across a wide range of forces following maximal effort fatiguing contractions

**DOI:** 10.1101/2023.04.26.538396

**Authors:** Emily J. Brotherton, Surendran Sabapathy, Saman Heshmat, Justin J. Kavanagh

## Abstract

People with Multiple Sclerosis typically exhibit greater levels of exercise-induced fatigue compared to healthy individuals. However, it is unknown if voluntary muscle activation is affected over a range of contraction forces in people with MS who have exercise-induced fatigue. The purpose of this study was to use transcranial magnetic stimulation (TMS) and electrical muscle stimulation to examine muscle activation during exercise-induced fatigue. Ten people with relapsing-remitting MS (39 ± 7 years) and 10 healthy controls (40 ± 5 years) performed elbow flexions at 25%, 50%, 75%, 90%, and 100% MVC while electromyography (EMG) of the biceps brachii was recorded. Sustained elbow flexion MVCs were then performed until force declined to 60% of baseline MVC, and the target contraction intensities of 25%, 50%, 75%, 90%, and 100% MVC were examined again. The Fatigue Severity Scale was higher for the MS group (*P* < 0.01). Exercise-induced fatigue caused a reduction in biceps EMG amplitude for the MS group across all contraction intensities (*P* < 0.01), which was not aligned with changes in MEP amplitude (*P* = 0.25). Exercise-induced fatigue reduced motor cortical voluntary activation in the MS group across all contraction intensities (*P* < 0.01), as well as increased MS time-to-peak force (*P <* 0.01) and half relaxation time for TMS evoked twitches (*P* = 0.03). These findings provide evidence that MS-related fatigability during maximal contractions is due to the inability for the motor cortex to drive the muscle, with possible contributions from altered contractile properties in the MS muscle.

**NEW & NOTEWORTHY:** We use transcranial magnetic stimulation to demonstrate that people with relapsing-remitting Multiple Sclerosis (MS) have a reduced ability to activate muscle following maximal effort fatiguing contractions. Although our MS participants reported greater symptoms of fatigue via the Fatigue Severity Scale, their reduced ability to activate muscle was more associated with the duration of disease.

## INTRODUCTION

Multiple sclerosis (MS) is an autoimmune neurodegenerative disorder that is characterised by progressive demyelination and subsequent neuronal loss within the central nervous system (CNS). For many people with MS, fatigue is the most debilitating symptom of the disorder, with up to 80% of people experiencing levels of fatigue that reduce their quality of life at some point in the disease course (1). The physiology underlying MS fatigue is complex, so steps are being made by MS researchers to operate within a framework where MS fatigue, and exercise-related fatiguability, are more clearly defined (2, 3). This contemporary approach highlights that feelings of fatigue in people with MS (state fatigue) should be differentiated from retrospective estimates of work capacity (perceived fatigability), and the ability to perform an exercise-related task (performance fatiguability) (2–4). Thus, a detailed approach to the measurement, and analysis, of fatigue-related data is critical for advancing our understanding of how physical activity may impact people with MS.

People with MS generally exhibit greater reductions in maximum voluntary force production for muscles of the upper limb (5–8) and the lower limb (9–11) during tasks that cause exercise-induced fatigue. We recently used transcranial magnetic stimulation (TMS) and motor nerve stimulation to examine performance fatiguability during prolonged low-intensity contractions of the elbow flexors (15% of maximal voluntary contraction [MVC] for 26 min). In doing so, we revealed that superimposed twitch responses, and hence voluntary activation of muscle, was compromised in relapsing-remitting MS compared to healthy controls (12). Not surprisingly, a strong association existed between higher scores in the clinically validated fatigue severity scale (FSS) and reductions in voluntary activation of muscle following prolonged low-intensity contractions in people with MS. Given that most differences were restricted to TMS activation of the motor cortex, it was apparent that people with MS had reduced output from the motor cortex while performing prolonged low-intensity contractions.

Although this work provided valuable insight to the MS motor system, it is well known that exercise-induced fatigue is dependent on several factors such as the duration and intensity of contraction (13). Thus, fatiguability for a low-intensity contraction may not reflect fatiguability for high-intensity contractions. In healthy individuals, fatigue-inducing maximal effort isometric contractions impairs the ability of the CNS to drive the muscle. For isometric elbow flexions, the performance of a maximal contraction that reduces force to 60% of an unfatigued MVC can reduce the ability of activate the muscle by up to 14% (14). Given that people with MS can have a reduced ability to activate muscle even before undertaking exercise (12), it is likely that fatiguing maximal contractions will inflict a significant challenge on the MS motor system (15). Functional magnetic resonance imaging has revealed that intracortical activity is lower in MS than healthy individuals during and following maximal fatiguing contractions (7). This potentially indicates that people with MS may not be able to increase synaptic input to motor cortical neurons experiencing a reduction in excitability due to the fatiguing task, which causes an inability to voluntarily activate muscle (3, 7). Nonetheless, the relationship between voluntary force and voluntary activation in people with MS performing very strong contractions is unknown.

The purpose of this study was to examine performance fatigability in people with MS associated with sustained fatigue-inducing maximal contractions of the elbow flexors. Measurements of voluntary activation were derived from twitch responses evoked by TMS and motor nerve stimulation during elbow flexions of 25%, 50%, 75%, 90% and 100% of MVC. These range of contraction intensities were examined under baseline condition (no intervention) and following a maximal effort sustained elbow flexion task that generated exercise-induced fatigue in the elbow flexors. We hypothesised that the MS group would exhibit greater declines in TMS-evoked voluntary activation and biceps EMG only following the performance of a maximal effort sustained elbow flexion. We hypothesised that these declines in voluntary activation and EMG would not be specific to a particular level of muscle contraction, but instead would be present for any contraction intensity performed after the fatigue-inducing task.

## MATERIALS AND METHOD

### Experimental design and ethical approval

The current study was a human, case-controlled, design, where all participants attended 2 testing sessions. One testing session included an exercise intervention whereas the other session had no intervention. The order of testing sessions was counter balanced and each session was separated by 2 weeks. Each participant provided written informed consent prior to undertaking testing. All experimental procedures were approved by the Griffith University Human Research Ethics Committee (reference number: 2020/338) and conducted according to the standards set by the Declaration of Helsinki.

### Participants

Ten people with MS (39 ± 7 years, 6 female) and 10 age-, gender-, and physical activity-matched healthy controls (40 ± 5 years, 6 female) were recruited into the study. MS participants required a confirmed diagnosis from a neurologist and must have been physically active at least 2-3 times per week for the previous 6 months. People with MS were ineligible for the study if they had a MS relapse within 30 days of testing, peripheral nerve pathology, joint of muscle pain in arms or legs, or any neurological comorbidities. All participants completed a medical history questionnaire prior to testing to ensure that they had no contraindications to transcranial magnetic stimulation and electrical stimulation. Trait levels of perceived fatigability were obtained from all participants prior to testing using the Fatigue Severity Scale (FSS). Medication schedules for the MS group were maintained prior to testing. All participants were asked to refrain from any form of CNS stimulant or depressant such as caffeine, alcohol, or moderate-to-high intensity exercise for twelve hours before testing. A priori statistical analysis using effect sizes based off group differences seen in neuromuscular activation (12) and was conducted in G*Power (v3.1.9.7) to determine the adequate sample size. To achieve an alpha of 0.05 and a power of 0.8 for 2 groups with a repeated measures ANOVA, 20 subjects were required to participate in the study.

### Electromyography and force

Participants were seated with the elbow of their dominant arm in 90 degrees of flexion and fixed in an aluminium device by two non-compliant straps (Figure 1A). Elbow flexion force was measured by a calibrated PT4000 S-Type tension and compression load cell (200kg, PT Ltd., New Zealand) with a 1.1 kN range and full-scale output of 3 mV/V. Force data was converted to torque for each participant using a lever arm based on the distance between the lateral epicondyle and the point of force application at the wrist. Surface EMG was obtained from the biceps and triceps brachii in the dominant arm by using two Ag/AgCl electrodes (Kendall ARBO, 24 mm) over each muscle in a muscle-tendon configuration. All surface EMG signals were amplified (x300) and bandpass filtered (10-1000 Hz) using a 2nd order Butterworth filter (CED 1902, Cambridge Electronic Design Ltd., UK). All EMG and force data was sampled at 2000 Hz using a Power 1401 data acquisition interface with Spike2 software (version 7.02, Cambridge Electronic Design Ltd., UK).

**Figure 1.**
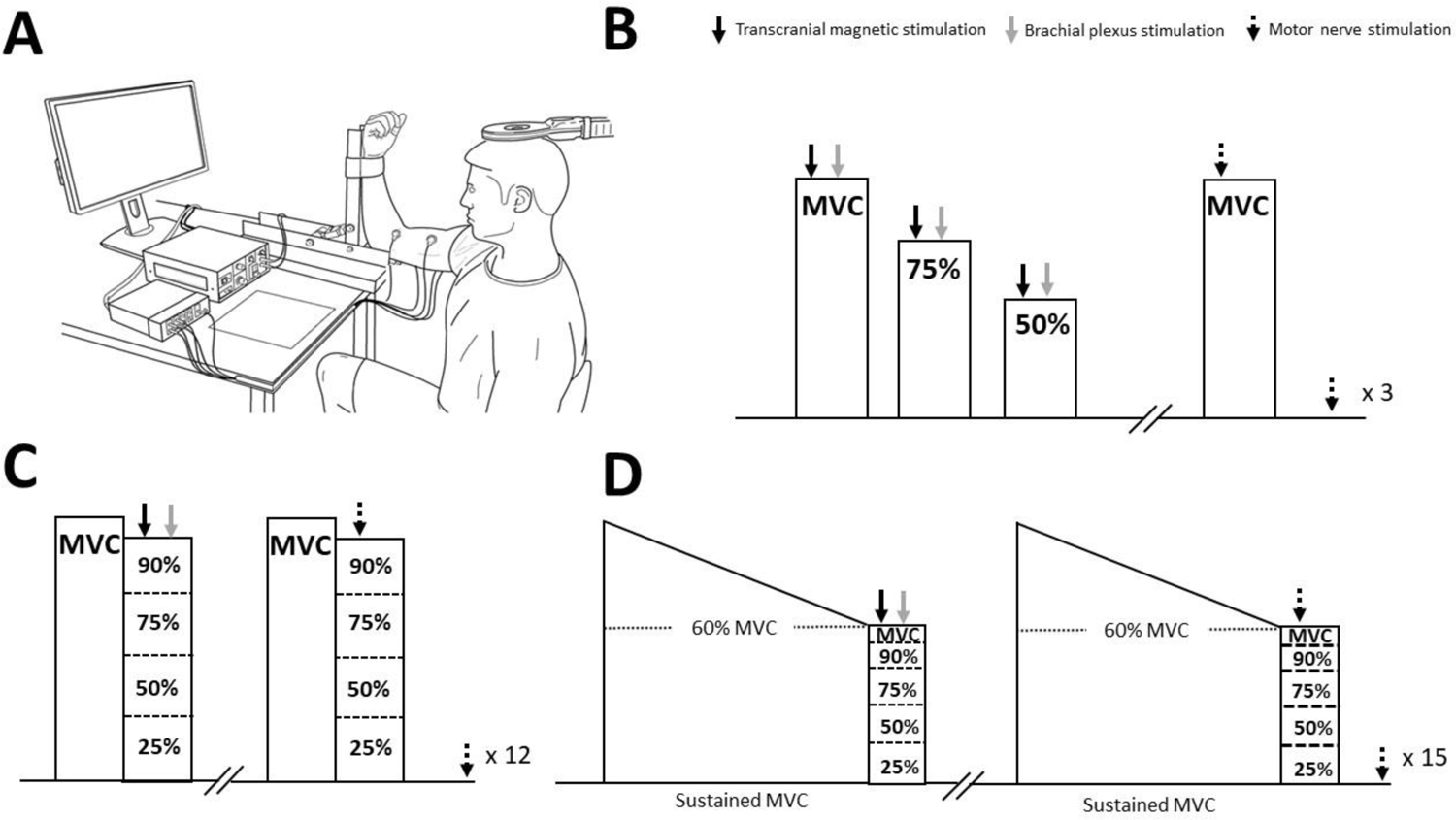
Experimental set-up and protocol. (A) Elbow flexion torque and biceps brachii EMG were measured during two contraction protocols performed in separate sessions. (B) Prior to each protocol, baseline data was collected to establish MVC, 75% MVC, and 50% MVC for that session to estimate TMS-evoked resting twitches. TMS was applied to the motor cortex and electrical stimulation was applied to the brachial plexus 2 s later for each contraction intensity. Baseline MVCs were also performed where a motor nerve stimulation was delivered to the biceps brachii during the contraction and another motor nerve stimulation was delivered ∼3 s following the contraction in the potentiated resting muscle. (C) During the session without the intervention, participants performed a brief (∼2 s) maximal elbow flexion to potentiate the muscle before immediately continuing with a 25% MVC, 50% MVC, 75% MVC, 90% MVC, or MVC. For each contraction intensity, TMS and brachial plexus stimulation was delivered during contractions, whereas motor nerve stimulation was delivered during and follow contractions. (D) For the session with the exercise intervention, sustained maximal elbow flexions were performed until force reduced to 60% of baseline MVC for that session. Similar to the first session, TMS, brachial plexus, and motor nerve simulation data was obtained for contraction intensities of 25% MVC, 50% MVC, 75% MVC, 90% MVC, and MVC.

### Brachial plexus stimulation

Supramaximal electrical stimuli were applied to the brachial plexus (100 μs pulse width, DS7AH constant current Digitimer Ltd., UK) to elicit a compound action muscle potential (M_max_) in the biceps and triceps brachii. A surface cathode was placed over the supraclavicular fossa (Erb’s point) with the positioning determined by using a motor nerve stimulating pen (at a stimulus intensity of 20 mA), and a surface anode was placed over the acromion. For both testing sessions, the stimulus intensity was progressively increased until a peak in the amplitude in the Mmax of both the biceps and triceps brachii was reached. The stimulation intensity was then fixed at 30% above the intensity required to elicit a maximal biceps and triceps M_max_, which was not significantly different between groups (MS: 171.4 ± 53.1 mA; Controls: 133.7 mA ± 29.3 mA) or testing sessions (Session 1: 151.4 mA ± 47.0 mA; Session 2: 154.4 mA ± 48.1 mA).

### Motor cortical stimulation

Transcranial magnetic stimulation (TMS; Magstim 200^2^ Co., UK) using a circular coil positioned over the vertex of the motor cortex (90 mm Remote Coil, MagStim Co., UK) was used to elicit motor evoked potentials (MEPs) in the biceps and triceps brachii muscles of the test limb. The orientation of the coil was positioned to activate the neural pathways in the left motor cortex to innervate the biceps brachii muscle of the participant’s right arm. The stimulator output of the Magstim was set at the lowest intensity that produced a biceps MEP > 60% of the biceps M_max_ and a triceps MEP < 20% of triceps M_max_ during brief unfatigued MVCs. This ensured that the TMS pulse evoked discharge of the largest proportion of motoneurons in the target muscle motoneuron pool (biceps brachii) whilst minimising the discharge of motoneurons in the motoneuron pool of the antagonist muscle (triceps brachii). The stimulator output was similar between groups (MS: 71.5% ± 3.2%; Controls: 70.5% ± 3.5%, *P* = 0.535) and testing sessions (Session 1: 70.5% ± 3.3%; Session 2: 71.0% ± 3.4%, *P* = 0.818).

### Motor nerve stimulation

Supramaximal electrical stimuli (100 μs pulse width) were delivered via a constant current stimulator (DS7AH, Digitimer Ltd., UK) to the intramuscular nerve fibres innervating the elbow flexors of the dominant arm. A surface cathode was positioned over the muscle belly and a surface anode placed over the distal tendon of the biceps brachii. A maximal resting twitch was obtained in the relaxed muscle by gradually increasing the stimulus intensity until no further increases in peak elbow flexion torque were achieved. The stimulation intensity was then set at 20% above the intensity required to elicit a maximal resting twitch and was similar between groups (MS: 135.6 mA ± 23.0 mA; Controls: 117.9 mA ± 35.2 mA, *P* = 0.175) and testing sessions (Session 1: 125.6 mA ± 32.1 mA; Session 2: 132.0 mA ± 33.1 mA, *P* = 0.643).

### Contraction protocols

Participants visited the lab on two occasions. Both testing sessions measured voluntary activation of the elbow flexors across a range of contraction intensities, however one of the sessions included a fatigue-inducing exercise intervention that consisted of a sustained MVC that preceded the range of contraction intensities. During each session, baseline measurements were performed where participants performed 6 brief (∼4 s) control MVCs (Figure 1B). Motor cortex and brachial plexus stimulations were delivered during 3 MVCs. Motor nerve stimulation was delivered during the other 3 MVCs, where an additional single stimulus was delivered ∼3 s after contraction at rest to evoke a potentiated resting twitch. Motor cortex and brachial plexus stimulations were also delivered during elbow flexions of 75% MVC and 50% MVC so that linear regression could be used to provide an estimate of resting twitch for TMS data (see data analysis). A rest period of at least 3 min was given between each contraction.

### Protocol without an intervention

The highest peak torque of the control MVCs was used to calculate submaximal target forces of 25%, 50%, 75% and 90% MVC. Participants then completed 24 brief (∼2 s) MVCs to potentiate the muscle, with each MVC immediately followed by a submaximal contraction in random order (Figure 1C). Six contractions were performed for each of the 4 submaximal target forces. During 3 of the 6 submaximal contractions, stimulation of the motor cortex and brachial plexus were delivered within the same contraction. For the other 3 submaximal contractions motor nerve stimulation was delivered during the contraction, where an additional single stimulus was delivered ∼3 s after contraction to evoke a potentiated resting twitch. A rest period of at least 3 min was given between each contraction, where participants were allowed additional time for recovery if they requested it.

### Protocol with a fatigue-inducing intervention

Participants completed 30 sustained maximal elbow flexion contractions until force declined to 60% of baseline MVC collected for that session (Figure 1D). Twenty-four of the sustained MVCs were immediately followed (< 1s) by a submaximal contraction of 25%, 50%, 75%, or 90% of the new ‘fatigued’ MVC, with the remaining six contractions being a standalone sustained MVC. Six contractions were performed for each of the target forces in random order. During three of the six contractions at the same intensity, stimulation of the motor cortex and brachial plexus was applied with motor nerve stimulation delivered during the other three submaximal contractions. When motor nerve stimulation was delivered during the submaximal contraction, an additional single stimulus was delivered ∼3 s after contraction at rest to evoke a potentiated resting twitch. Participants determined the rest period between each of the sustained MVC contraction tasks so that they were comfortable performing the next contraction in the session.

### Data analysis

All torque and EMG data were collected in Spike2 (version 7.02, Cambridge Electronic Design) and processed as waveform averages offline using Signal v6 software (Cambridge Electronic Design Ltd., UK). Peak amplitude of torque was collected from MVCs, and root mean square of the EMG (EMG_RMS_) of the biceps brachii and triceps brachii were measured during a 200 ms window prior to the stimulus artefact. The peak-to-peak amplitude of superimposed and resting twitch data was measured from the stimulus artefact to the peak of the increased torque evoked by TMS and motor nerve stimulation. Motor cortical and motor nerve voluntary activation (%) was calculated via the equation: (1 – superimposed twitch/resting twitch) x 100. For motor nerve voluntary activation, the resting twitch was measured from the change in torque amplitude from stimulation in the potentiated muscle at rest (16). For motor cortical voluntary activation, the resting twitch torque was estimated from the linear relationship of superimposed twitch responses at 100 %, 90%, 75% and 50 % MVC for unfatigued contractions and 100 %, 90%, 75% and 50% of the fatigued MVC for fatigued contractions. Data for contraction intensities of 25% MVC were not analysed due to non-linearity in twitch responses (14). A linear regression of twitch responses plotted against the corresponding voluntary torque was used to calculate the y-intercept, which represented the estimated resting twitch (17). MEP and M_max_ area for the biceps was calculated between vertical cursors that included all phases of the waveform. MEPs evoked from TMS were normalised to the corresponding M_max_ elicited by brachial plexus stimulation 2 s later to account for any activity-dependant changes in muscle fibre properties.

### Statistical analysis

All analyses were performed in R using RStudio (version 4.1.1; R Foundation for Statistical Computing, Vienna, Austria). Student T-tests were used to determine groups differences in participant characteristics and baseline neurophysiological variables. Mauchly’s test was used to identify data that violated sphericity, and when significant, Greenhouse-Geisser corrections were applied. Linear mixed effect models were used to examine the main effects of group (MS, Control), contraction condition (no intervention, fatigue-inducing exercise intervention) and contraction intensity (25%, 50%, 75%, 90% and 100% MVC) on outcome parameters (biceps brachii RMS EMG, biceps brachii MEP area, TMS superimposed twitch torque, time-to-peak for TMS superimposed twitch torque, TMS voluntary activation, motor nerve superimposed twitch torque, time-to-peak for MP superimposed twitch torque, motor nerve resting twitch torque and motor nerve voluntary activation) using *nlme* package in R. Models were created using iteratively adding predictor variables or interaction effects and determined by model comparison with the lowest Akaike Information Criterion (AIC) and statistical significance (*P* < 0.05) performed by an ANOVA accepted as the most appropriate model. The best fitting model included group, contraction intensity and FSS as fixed variables with a random intercept for each subject. Interaction effects between group and time were also examined. Correlation analyses were also performed for each group to determine if trait levels of perceived fatigability (FSS) and disease duration were correlated with neurophysiological measurements at baseline, following no intervention and following the exercise intervention. An alpha level of 0.05 was used to indicate statistical significance for all statistical tests. Data summarised in the text are presented as mean and standard deviation.

## RESULTS

### Participant characteristics and baseline variables

The MS group had an average disease duration of 7 years, and consequently reported significantly higher ratings for the fatigue severity scale compared to the control group in both sessions (Table 1). There were no other group differences for participant characteristics or baseline measurements.

**Table 1.** Participant characteristics and baseline measurements

|  | Multiple Sclerosis<br>n = 10 | Healthy controls<br>n = 10 |
| --- | --- | --- |
| *Characteristic* |  |  |
| Age (yr) | 39 ± 7 | 40 ± 5 |
| Weight (kg) | 76.9 ± 4.4 | 69.9 ± 11.7 |
| Height (m) | 1.77 ± 0.10 | 1.70 ± 0.06 |
| Disease duration (yr) | 7.6 ± 4.3 | N/A |
| *Fatigue severity scale* |  |  |
| Session 1 | 4.4 ± 1.6* | 2.9 ± 0.7 |
| Session 2 | 4.7 ± 1.5* | 2.8 ± 0.6 |
| *Baseline measurements* |  |  |
| MVC torque (N.m) | 52.45 ± 20.40 | 53.87 ± 15.38 |
| MVC biceps brachii RMS EMG (mV) | 0.596 ± 0.355 | 0.820 ± 0.378 |
| Motor point resting twitch (N.m) | 9.82 ± 3.54 | 10.15 ± 3.50 |
| TMS estimated resting twitch (N.m) | 11.55 ± 4.88 | 12.44 ± 4.35 |
| Motor point level of voluntary activation (%) | 97.1 ± 2.4 | 97.6 ± 1.5 |
| TMS level of voluntary activation (%) | 95.6 ± 2.4 | 97.9 ± 1.5 |

Maximal voluntary contraction, MVC; Transcranial magnetic stimulation, TMS; Asterisks denotes a significant difference between groups ( $p < 0.01$ )

### EMG amplitude and MEPs during voluntary contractions

The amplitude of EMG and motor evoked potentials were measured in the biceps brachii during the performance of elbow flexions of varying contractions intensities (25% MVC to 100% MVC). Biceps EMG amplitude linearly increased with the level of force that was being produced (main effect of contraction; F_4,71_ = 46.163, *P* < 0.0001, Figure 2A). This increase in EMG activity did not differ between groups, as there was no main effect detected for group (F_1,18_ = 2.183, *P* = 0.512), and no group by contraction intensity identified for biceps EMG amplitude (F_4,64_ = 0.109, *P* = 0.979). The relationship between EMG and force tended to be more curve linear following the performance of the sustained maximal effort contraction, where control group maximum EMG amplitude (i.e. during the during MVC) was 0.99 ± 0.40 mV and 0.98 ± 0.43 mV with and without the exercise intervention respectively (Figure 2B). This contrasted the MS group where maximum EMG amplitude was 0.52 ± 0.31mV and 0.75 ± 0.52 mV with and without the exercise intervention respectively. Following sustained maximal contractions, main effects for contraction intensity (F_4,75_ = 35.78, *P* < 0.0001) and group (F_1,18_ = 9.526, *P* = 0.006) were identified, where the MS group exhibited reduced EMG amplitude during elbow flexions. A group by contraction interaction was detected (F_4,71_ = 4.833, *P* = 0.002) for biceps brachii EMG during elbow flexions.

**Figure 2.**
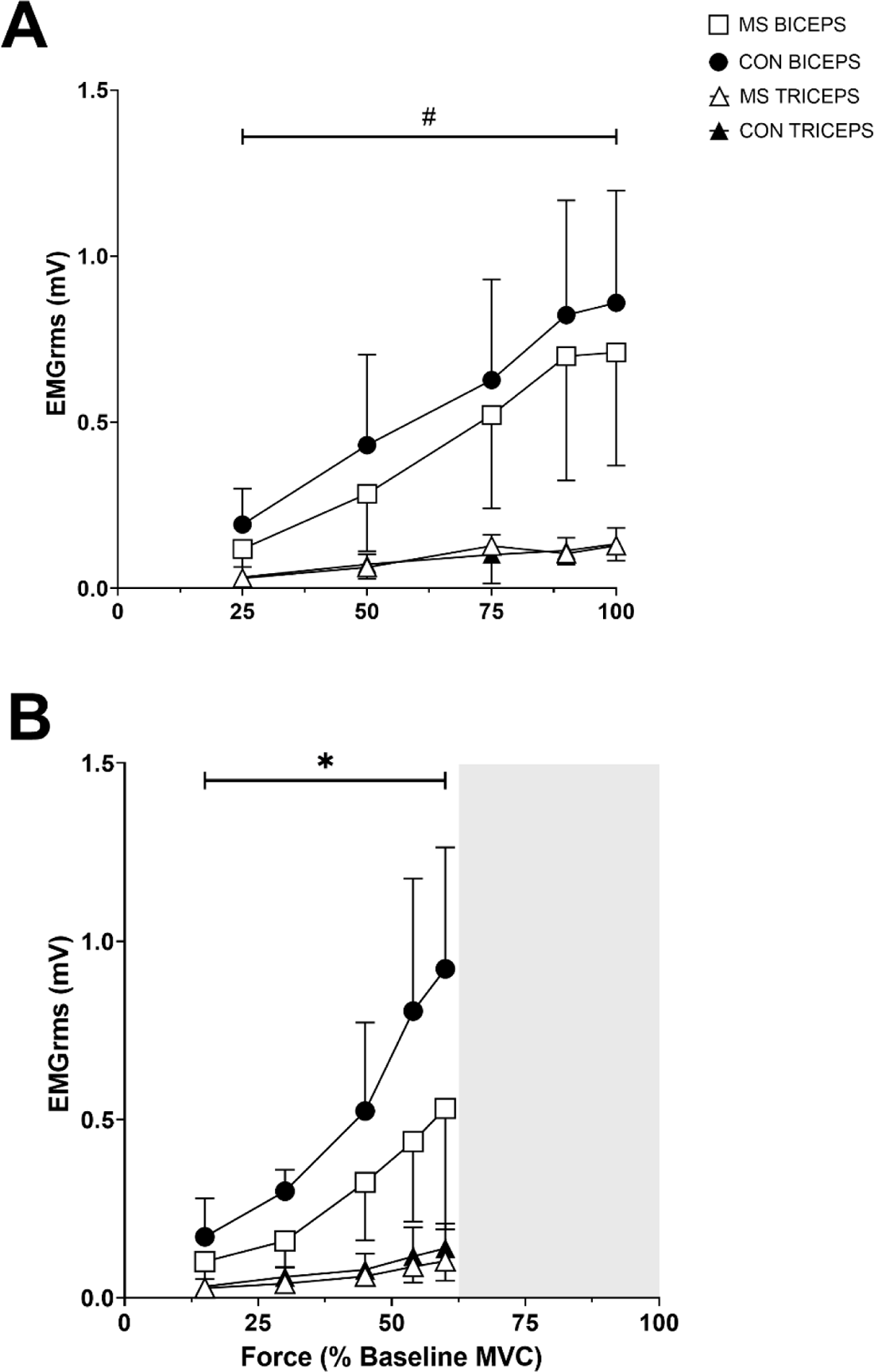
Biceps brachii and triceps brachii EMG amplitude. RMS EMG amplitude for the biceps brachii and triceps for the session without an intervention (A) and the session with the exercise intervention (B). Biceps and triceps RMS EMG is expressed as absolute values that were obtained during contraction intensities of 25%, 50%, 75%, 90% and 100% MVC. Although all forces are presented as percentages of MVC, the contraction intensities associated with the fatigue-inducing task in panel B are based on the new ‘fatigued’ MVC where force has substantially reduced from baseline (i.e 60% of the baseline MVC). Thus, the shaded area on the lower panel highlights that the fatigued contraction intensities are inherently lower than baseline trials. Hash symbol indicates a main effect of contraction intensity, and an asterisk indicates a main effect of group. All data are presented as mean ± SD (MS group *n* = 10, Control group *n* = 10).

For both groups the normalised MEP increased from 25% MVC to 50% MVC before declining in amplitude as the participants approached MVC (Figure 3A). Although there was a main effect of contraction intensity (F_4,71_ = 8.830, *P* < 0.001), there was no main effect of group (F_1,18_ = 1.3745, *P* = 0.256), or group by contraction intensity interaction (F_4,67_ = 1.489, *P* = 0.216) for biceps brachii MEP during elbow flexions. The sustained maximal contraction altered the relationship between MEP and force, as the normalised MEP increased from 25% to 75% of the fatigued MVC in controls and from 25% to 90% of the fatigued MVC in MS, before declining in amplitude towards MVC (main effect of contraction intensity, F_4,75_ = 9.466, *P* < 0.001, Figure 3B). However, MEPs following the sustained contractions were not altered with MS, as no main effect of group (F_1,18_ = 2.221, *P* = 0.154) and no group by contraction intensity interaction (F_4,71_ = 1.302, *P* = 0.278) was detected for biceps brachii MEP during elbow flexions.

**Figure 3.**
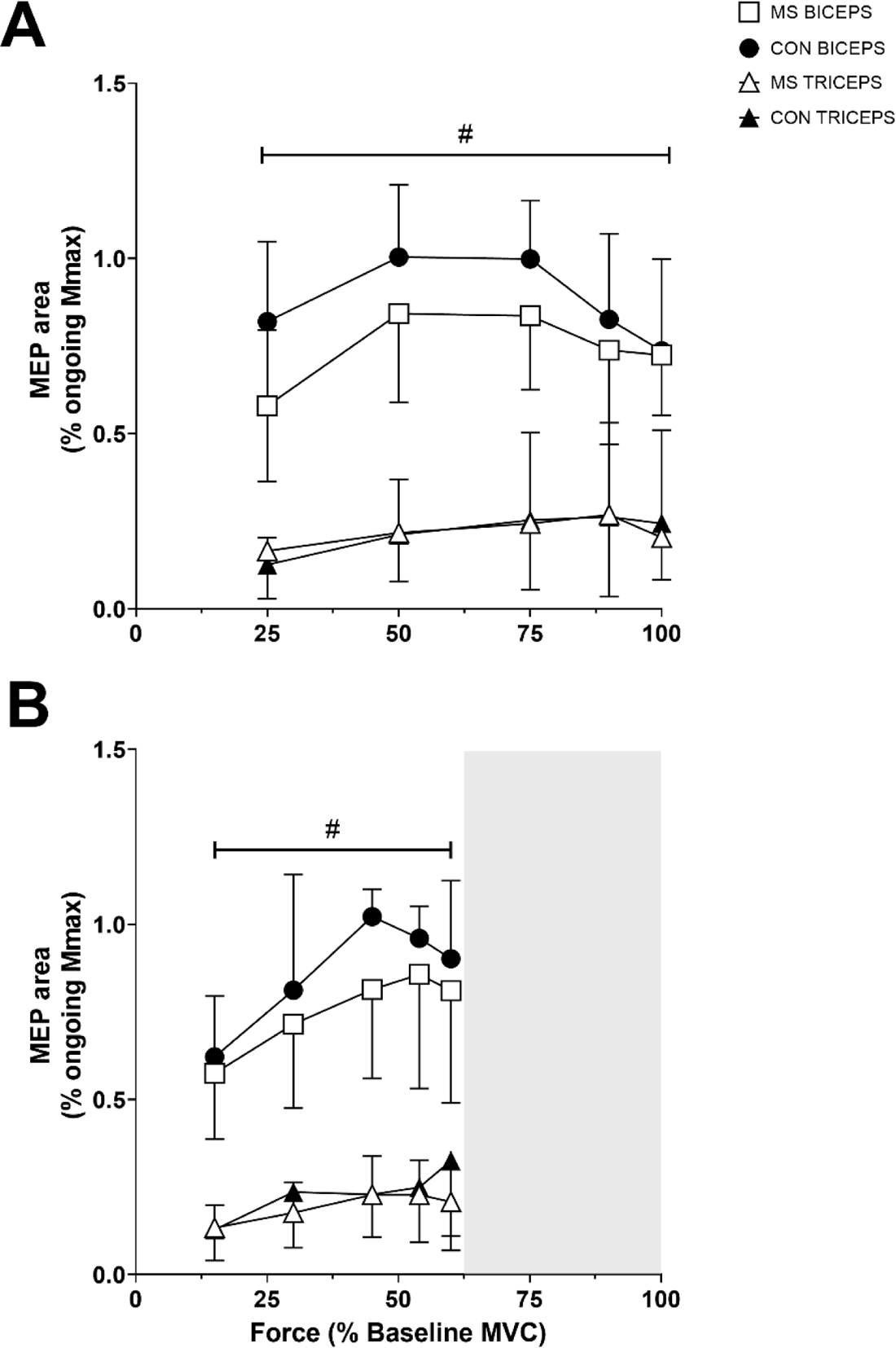
Motor evoked potentials to motor cortical stimulation during varying contraction intensities. MEP area for biceps brachii for the session without an intervention (A) and the session with the exercise intervention (B). MEP is expressed as a percentage of the biceps maximal compound action potential (M_max_) area that was obtained during contraction intensities of 25%, 50%, 75%, 90% and 100% MVC. The shaded area on the lower panel highlights that the fatigued contraction intensities during the exercise intervention are presented normalised to the baseline MVC. Hash symbol indicates a main effect of contraction intensity (*P* < 0.001). All data are presented as mean ± SD (MS group *n* = 10, Control group *n* = 10).

### Twitch responses to motor cortical and motor nerve stimulation

Twitch forces evoked by motor cortical and motor nerve stimulation were measured in the elbow flexors across varying contractions intensities following brief maximal and sustained maximal contractions in both MS and controls. Figure 4 shows the average waveform of twitch forces evoked torque evoked in the elbow flexors across all contraction intensities in one person with MS and one healthy age-matched control.

**Figure 4.**
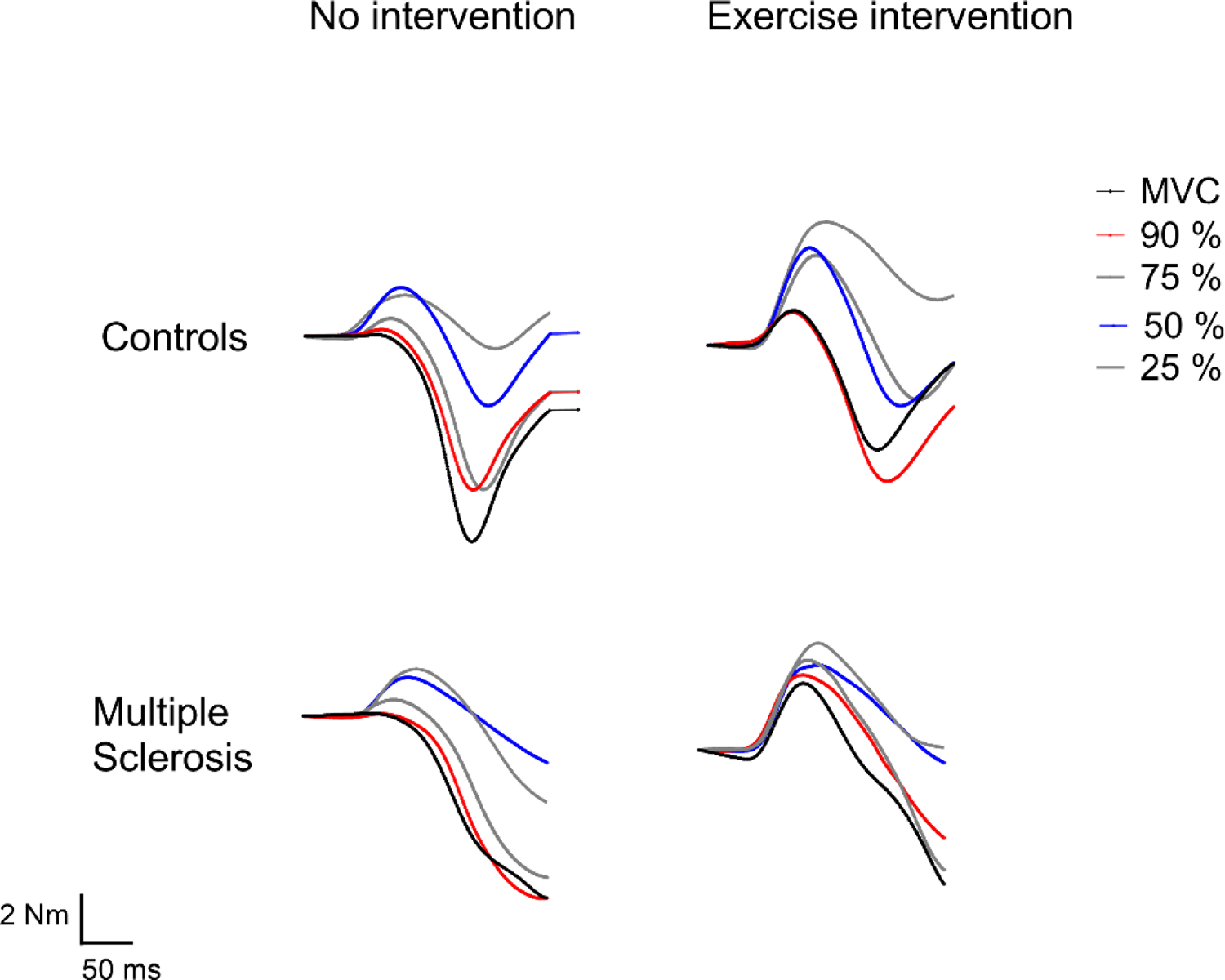
Motor cortical and motor nerve evoked twitches following brief and sustained maximal contractions. Averaged waveforms of twitch forces evoked in the elbow flexors by motor cortical and motor nerve stimulation during contraction intensities from 25% to 100% maximum voluntary contraction (MVC) from one person with MS (age: 41, years with MS: 5, FSS: 4.5) and one age-matched control. Superimposed twitches are presented for each contraction intensity following the no intervention and the exercise intervention protocol.

Motor cortical superimposed twitches progressively declined with increased contraction intensity for both the MS and control groups. Although there was a main effect of contraction intensity (F_4,70_ = 60.641, *P* < 0.001), there were no significant group differences (F_1,18_ = 1.624, *P* = 0.219), but a group by contraction intensity interaction (F_4,66_ = 3.532, *P* = 0.011), for motor cortical superimposed twitch amplitude (Table 2). Overall, the performance of the maximal effort contractions in the exercise intervention caused an enhancement of the motor cortical superimposed twitch amplitude at each contraction intensity for each group. Similar to brief maximal contractions, a main effect of contraction intensity (F_4,75_ = 25.814, *P* < 0.001) and group by contraction intensity interaction (F_4,71_ = 5.790, *P* < 0.001) was detected without a main effect of group (F_1,18_ = 0.124, *P* = 0.728) for motor cortical superimposed twitches following the sustained maximal contraction in the exercise intervention.

**Table 2.** Superimposed twitch amplitudes evoked by transcranial magnetic stimulation and motor nerve stimulation

|  | Motor cortical |  | Motor nerve |  |
| --- | --- | --- | --- | --- |
| *No intervention* | MS | CON | MS | CON |
| **Rest** | 12.5 ± 4.5 | 11.1 ± 3.8 | 10.3 ± 3.5 | 9.6 ± 3.7 |
| **25% MVC** | 5.1 ± 1.8 | 7.1 ± 2.6 | 6.2 ± 1.6 | 5.6 ± 1.7 |
| **50% MVC** | 5.6 ± 2.2 | 5.9 ± 1.6 | 3.3 ± 0.8 | 2.2 ± 0.5 |
| **75% MVC** | 2.6 ± 0.7 | 2.7 ± 1.1 | 1.5 ± 0.8 | 0.8 ± 0.4 |
| **90% MVC** | 1.0 ± 0.6 | 0.9 ± 0.5 | 0.6 ± 0.6 | 0.3 ± 0.2 |
| **100% MVC** | 0.5 ± 0.2 | 0.7 ± 0.5 | 0.3 ± 0.2 | 0.2 ± 0.1 |
| *Exercise intervention* | MS | CON | MS | CON |
| **Rest** | - | - | - | - |
| **25% MVC** | 5.9 ± 1.7 | 7.3 ± 3.5 | 6.6 ± 2.8 | 5.1 ± 3.0 |
| **50% MVC** | 6.8 ± 2.6 | 5.9 ± 2.5 | 4.1 ± 1.8 | 3.0 ± 2.1 |
| **75% MVC** | 5.5 ± 2.2 | 5.1 ± 1.9 | 2.6 ± 1.4 | 1.7 ± 1.3 |
| **90% MVC** | 4.7 ± 2.8 | 3.4 ± 1.5 | 1.8 ± 1.1 | 1.1 ± 0.6 |
| **100% MVC** | 3.9 ± 2.1 | 2.1 ± 1.1 | 1.2 ± 0.7 | 0.7 ± 0.6 |

Maximal voluntary contraction, MVC; All data are presented as mean ± SD (MS group $n = 10$ , Control group $n = 10$ ). Asterisks denotes a significant difference between groups ( $p < 0.05$ ).

Superimposed twitches evoked by motor nerve stimulation followed a similar pattern to those evoked via TMS. That is, motor nerve superimposed twitch amplitude progressively declined with increasing contraction intensity (F_4,67_ = 115.19, *P* < 0.001), and no significant group differences (F_1,18_ = 0.807, *P* = 0.389). However, there was no group by contraction intensity interaction (F_4,63_ = 0.914, *P* = 0.878) was detected during the contractions (Table 2). The performance of the maximal effort contractions in the exercise intervention once again caused an enhancement of the superimposed twitch amplitude at each contraction intensity, where a main effect of contraction intensity (F_4,75_ = 69.23, *P* < 0.001) was detected without a main effect of group (F_1,18_ = 1.129, *P* = 0.302) or a group by contraction intensity interaction (F_4,71_ = 0.296, *P* = 0.880) for motor nerve superimposed twitches.

### Time-to-peak force and half relaxation time following stimulation

Regardless of whether motor cortical or motor nerve stimulation was used to elicit twitch forces, the time-to-peak and half relaxation time progressively shortened from 25% MVC to MVC for both the MS group and the healthy controls (Table 3). No significant differences were detected between groups at any of the contraction intensities for the session without the intervention. However, following the exercise intervention, MS demonstrated a greater time-to-peak force elicited by both TMS (*P* = 0.005) and motor nerve stimulation (*P* = 0.016) at the fatigued MVC. Similarly, the half relaxation time elicited by TMS at the fatigued MVC (*P* = 0.03) was also significantly greater in MS compared to controls.

**Table 3.**
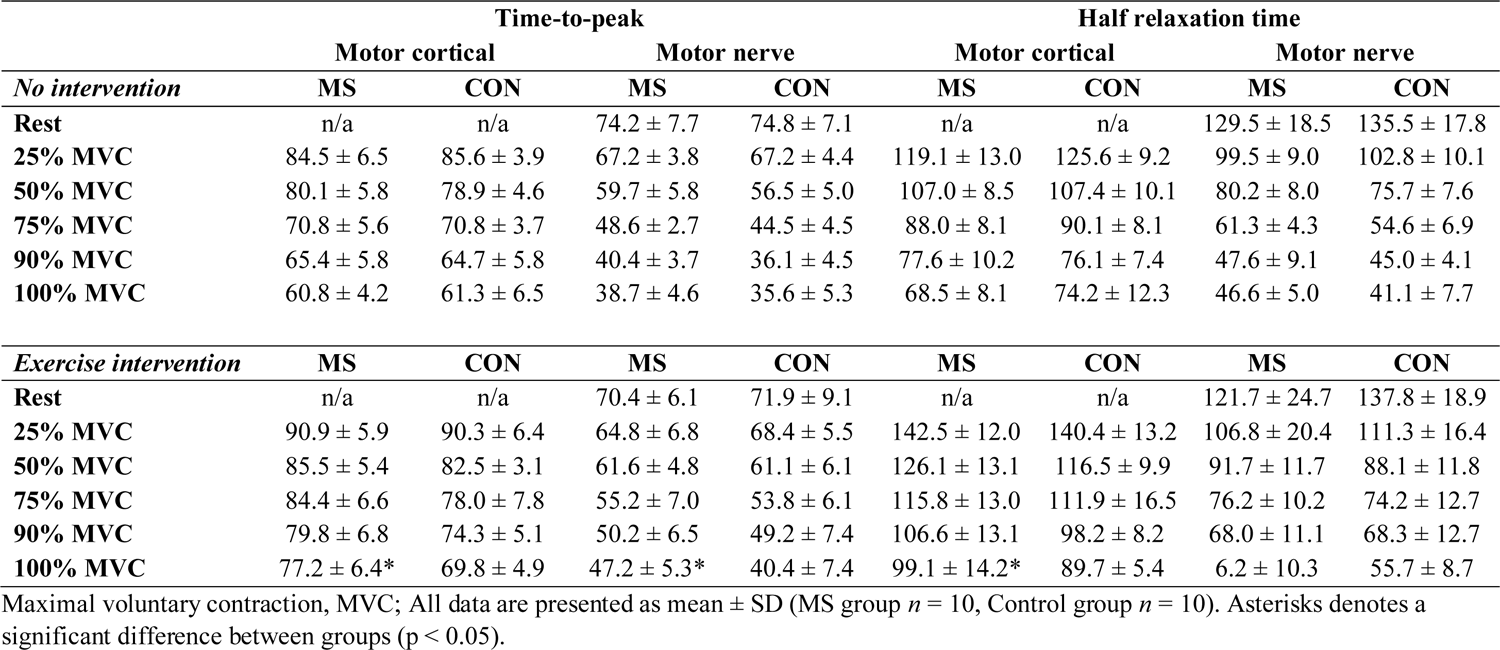
Time-to-peak and half relaxation time of superimposed twitch evoked by transcranial magnetic and motor nerve stimulation

### Voluntary muscle activation

A linear relationship existed for motor cortical voluntary activation in the MS group (r^2^ = 0.972 ± 0.031) and control group (r^2^ = 0.989 ± 0.015) following brief maximal contractions (Figure 5A) and in the MS group (r^2^ = 0.942 ± 0.031) and control group (r^2^ = 0.956 ± 0.035) following sustained maximal contractions (Figure 5B). This indicates that motor cortical voluntary activation closely reflected each group’s ability to generate elbow flexion force, with increases in voluntary activation matching increases in force (main effect of contraction intensity (F_4,70_ = 170.056, *P* < 0.001). Although there was no main effect of group for motor cortical voluntary activation (F_1,18_ = 1.013, *P* = 0.326), a group by contraction intensity interaction was detected (F_4,66_ = 4.501, *P* = 0.003). Performance of the sustained maximal contraction resulted in substantial changes in voluntary activation. While a main effect of contraction intensity was once again identified (F_4,75_ = 30.061 *P* < 0.001), there was also a main effect of group detected for motor cortical voluntary activation (F_1,18_ = 35.693, *P* < 0.001, Figure 5B) to accompany a group by contraction intensity interaction (F_4,71_ = 2.803, *P* = 0.032) for motor cortical voluntary activation. Voluntary activation for MS group was on average 22% lower for the corresponding force in the control group, with maximal voluntary activation in the MS group only reaching 67.26% ± 16.26% compared to 85.67% ± 4.90% in the control group.

**Figure 5.**
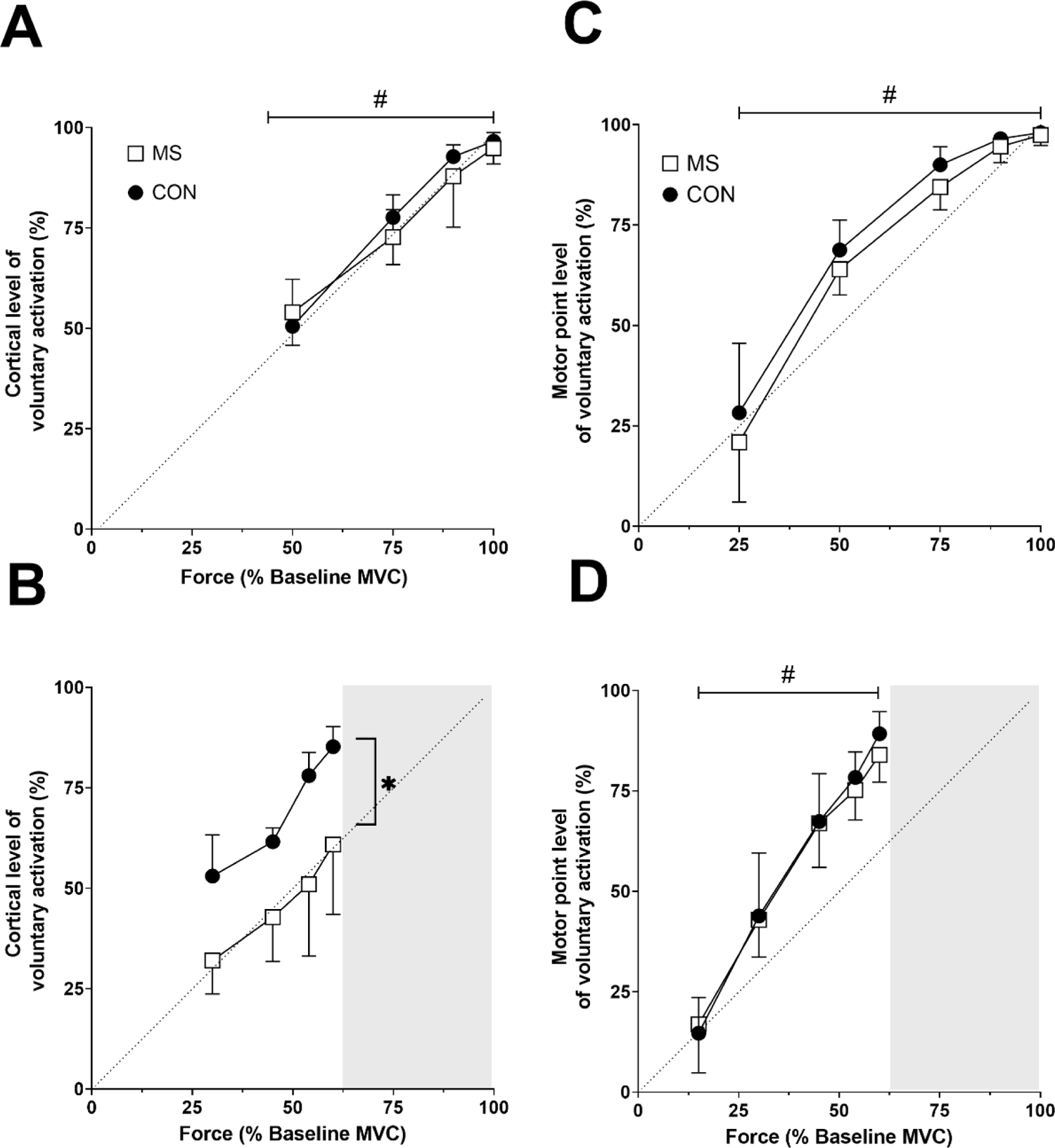
Voluntary muscle activation. Motor cortical voluntary activation (A and B), and motor nerve voluntary activation (C and D) was calculated for contraction intensities of 25%, 50%, 75%, 90% and 100% MVC. For motor cortical voluntary activation, data for 25% MVC were not analysed due to non-linearity in twitch responses. Although all forces are presented as percentages of MVC, the contraction intensities associated with the fatigue-inducing task in panels B and D are based on the new ‘fatigued’ MVC where force has substantially reduced from baseline. Hash symbol indicates a main effect of contraction intensity, and an asterisk indicates a main effect of group. All data are presented as mean ± SD (MS group *n* = 10, Control group *n* = 10).

A curvilinear relationship existed for motor nerve voluntary activation in both the MS group and controls following brief maximal contractions (range of r^2^ for MS: 0.83 – 0.91 range of r^2^ for controls: 0.87 - 0.98, however following sustained maximal contractions this relationship became more linear (range of r^2^ for MS: 0.96 - 0.99, range of r^2^ for controls: 0.95-0.99). Similar to motor cortical stimulation, increases voluntary activation calculated from motor nerve stimulation matched increases in force (main effect of contraction intensity; F_4,67_ = 270.054, *P* < 0.001). However, motor nerve voluntary activation did not detect any differences due to MS, as there was no main effect of group (F_1,18_ = 2.105, *P* = 0.164) or group by contraction intensity interaction (F_4,63_= 0.348, *P* = 0.845). Following brief maximal contractions, the maximal level of voluntary activation that could be achieved 88.13% ± 7.32% in the MS group which was similar to 88.54% ± 5.53 % in the control group (F_1,18_= 0.107, *P* = 0.746).

### Correlation between MS perceived fatigue severity, disease duration and voluntary muscle activation

There was no significant relationship identified between the FSS and any TMS or motor nerve stimulation measure of voluntary muscle activation (Table 4). However, when the duration of disease was used in bivariate correlation analysis, a strong and significant correlation was identified between disease duration and motor cortical voluntary activation (*R^2^* = 0.757; *P*= 0.011) following the sustained maximal contractions. A longer duration of MS corresponded to lower motor cortical voluntary activation after the fatigue-inducing elbow flexion task.

**Table 4.**
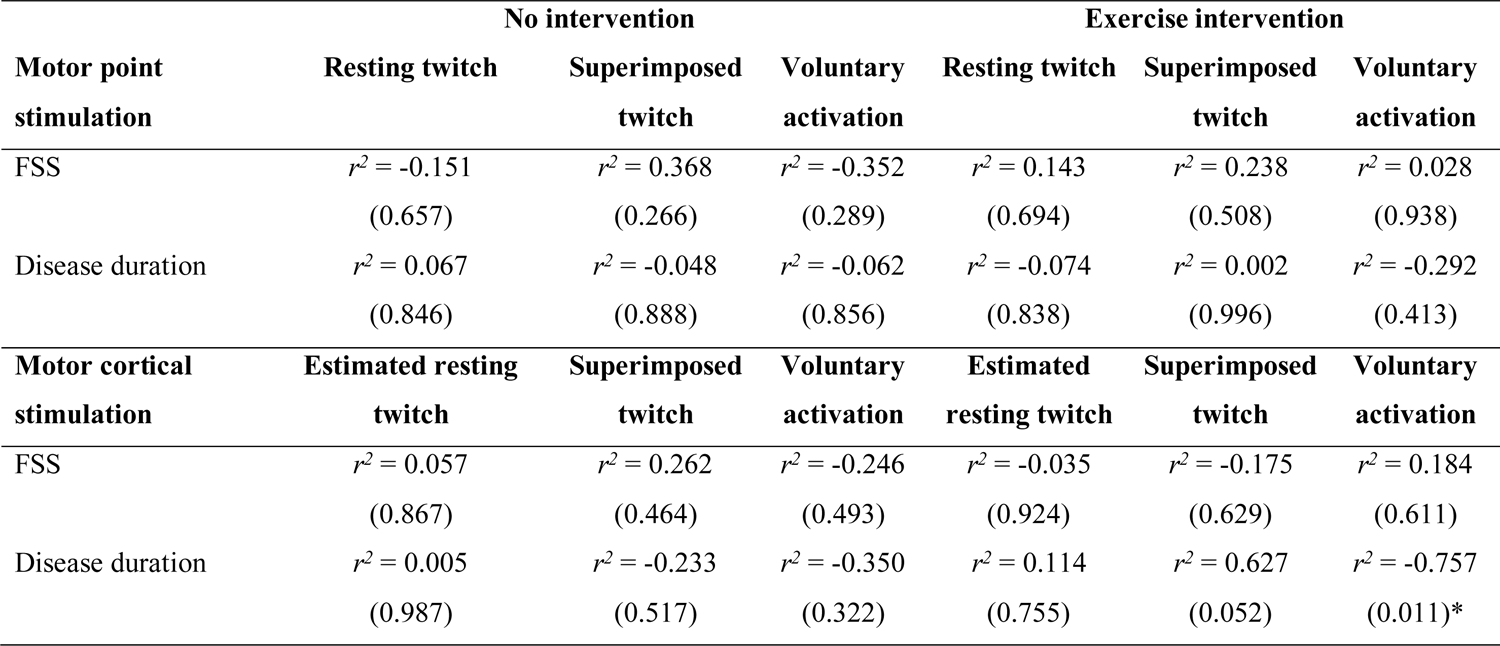
Correlation between MS disease severity and voluntary muscle activation

| **Motor point stimulation** | **No intervention** |  |  | **Exercise intervention** |  |  |
| --- | --- | --- | --- | --- | --- | --- |
|  | **Resting twitch** | **Superimposed twitch** | **Voluntary activation** | **Resting twitch** | **Superimposed twitch** | **Voluntary activation** |
| FSS | $r^2 = -0.151$<br>(0.657) | $r^2 = 0.368$<br>(0.266) | $r^2 = -0.352$<br>(0.289) | $r^2 = 0.143$<br>(0.694) | $r^2 = 0.238$<br>(0.508) | $r^2 = 0.028$<br>(0.938) |
| Disease duration | $r^2 = 0.067$<br>(0.846) | $r^2 = -0.048$<br>(0.888) | $r^2 = -0.062$<br>(0.856) | $r^2 = -0.074$<br>(0.838) | $r^2 = 0.002$<br>(0.996) | $r^2 = -0.292$<br>(0.413) |
| **Motor cortical stimulation** | **Estimated resting twitch** | **Superimposed twitch** | **Voluntary activation** | **Estimated resting twitch** | **Superimposed twitch** | **Voluntary activation** |
| FSS | $r^2 = 0.057$<br>(0.867) | $r^2 = 0.262$<br>(0.464) | $r^2 = -0.246$<br>(0.493) | $r^2 = -0.035$<br>(0.924) | $r^2 = -0.175$<br>(0.629) | $r^2 = 0.184$<br>(0.611) |
| Disease duration | $r^2 = 0.005$<br>(0.987) | $r^2 = -0.233$<br>(0.517) | $r^2 = -0.350$<br>(0.322) | $r^2 = 0.114$<br>(0.755) | $r^2 = 0.627$<br>(0.052) | $r^2 = -0.757$<br>(0.011)* |

## DISCUSSION

The purpose of this study was to examine performance fatigability in people with MS during sustained maximal fatiguing contractions of the elbow flexors. Measurements of voluntary activation were derived from TMS and motor nerve stimulation during a wide range of elbow flexion contraction intensities. Measurements were obtained in two testing sessions where one session employed a maximal effort sustained elbow flexion task that generated exercise-induced fatigue in the elbow flexors. The main findings of the study were that following the maximal sustained contractions the MS group had significantly lower biceps EMG amplitude, and voluntary activation measured via TMS, compared to healthy controls. Although there were distinct differences in subjective levels of fatigue between MS and controls (MS FSS > 4 compared to control FSS < 3), the disease duration was the best predictor of voluntary muscle activation following the fatigue inducing exercise intervention.

### Biceps brachii muscle activity and MEPs

EMG activity produced during voluntary contractions progressively increases with an increase in contraction intensity. This relationship is typically attributed to increases in motor unit recruitment and/or firing rate to maintain a force target (18), and once the upper limit for motor unit recruitment increases in force are predominantly due to increases in firing rate (19, 20). Therefore, it is not surprising that both groups demonstrated an increase in RMS EMG activity of the biceps brachii with an increase in contraction intensity following both normal muscle contraction and the exercise-induced fatiguing task. However, between group differences were only found for contractions following the fatiguing task where the MS group had significantly lower EMG amplitude following the sustained maximal contraction. Furthermore, the difference between groups became more apparent at higher contraction intensities, which suggests that the MS group had a greater inability to increase motor unit firing rate to match the demands of the contraction task once all motor units were recruited.

The findings in the current study support previous work where individuals with MS exhibited a reduction in biceps brachii EMG during a fatiguing contractions during a brief maximal contraction of the elbow flexors (12) and knee extensors (9). Interestingly, the mechanism underlying reduction in MS muscle activity may not be associated with excitability of neurons in the motor pathway, as there was no change in MEPs during the contractions performing with, or without, the exercise intervention. This aligns with previous investigations that report people with MS have similar corticospinal excitability to healthy individuals during fatiguing exercise (5, 12, 21, 22). Notably, previous studies of the upper limb showed that a fatiguing contraction (sustained MVC (5, 21) and sustained low-intensity (12)) has similar effect on the MEP area in both fatigued MS (FSS > 4) and healthy controls. Building on this, the recent works of Leodori et al. (22) demonstrated that fatigued people with MS and healthy controls have similar corticospinal excitability and corticospinal tract transmission during an intermittent fatigue index finger contractions. Thus, a weight of evidence suggests that neuronal excitability in motor pathways that directly activate muscle may not be the primary reason underlying greater performance fatiguability in MS.

### Voluntary muscle activation

Twitch responses to stimulation were used to calculate voluntary muscle activation, where superimposed twitch amplitude closely aligned with voluntary activation for both TMS and motor nerve stimulation. Surprisingly, the relationship between voluntary force and voluntary activation has not been quantified for people with MS. Thus, it is unknown if MS requires more motor cortical output to activate muscles when performing similar contraction intensities to healthy individuals. In healthy individuals, the relationship between voluntary force production and measures of voluntary activation of both the unfatigued and fatigued elbow flexors increases in a curve-linear manner when measured with motor nerve stimulation and a linear manner when measured with TMS (17). The current study provides evidence that for the rested (non-fatigued) elbow flexors of individuals with MS, the relationship between voluntary force production and measures of voluntary activation is comparable to healthy individuals when measured with motor nerve stimulation and TMS. However, following a sustained maximal elbow flexion contraction, people with MS demonstrated a reduction in voluntary activation measured by TMS across all contraction intensities. These findings suggest that performance fatigability experienced by individuals with MS during fatiguing contractions is due to the inability of the motor cortex to drive the muscle maximally.

The clinical course of MS is characterised by relapses and disease progression secondary to structural damage to both white and grey matter (23, 24). Functional cortical changes due to axonal demyelination and damage have been shown to result in altered cortical activation during exercise (5, 25–27). Moreover, previous functional magnetic resonance imaging (fMRI; 28, 29) and electroencephalogram (EEG; 22, 30) studies have shown that MS is associated with increased cortical activation during a submaximal task compared to controls, possibly, in an attempt conserve cortical output despite cortical damage. Thus, it is possible that the influence of MS disease duration on white/grey matter can cause substantial motor cortical deficits despite being physically active with mild disability. Our findings of a strong correlation between disease duration and motor cortical voluntary activation deficits further supports the notion that cortical dysfunction is associated with disease progression in MS.

The motor cortical impairments during fatiguing exercise in the current study align with findings from our previous work (12). This experiment revealed that people with MS have heightened performance fatiguability during a low intensity sustained elbow flexion task which is accompanied by greater reductions in neural drive to the muscle than healthy individuals. In contrast, Coates et al. found no differences in motor cortical voluntary activation detected between MS and healthy individuals following a fatiguing lower limb cycling task (9). However, these disparity in findings could be attributed to a number of factors, including the ability to assess TMS-related voluntary in upper and lower limb muscles (17, 31), different MS inclusion criteria for each study, and vastly different exercise requirements when performing a sustained isometric contraction compared to a dynamic cycling challenge. It is important to note that the voluntary activation findings of the current study must be interpreted in the context of relapsing-remitting MS, and findings may be applicable to other MS cohorts. The participants in the current had no recent history of relapse and ultimately were similar for many physiological processes to healthy controls when there was no exercise intervention. However, despite undertaking physical activity 2-3 times per week, our MS participants still appeared to be compromised during the exercise intervention, which highlights the inherent challenges that people with MS can face when performing vigorous exercise.

### Contractile properties of the muscle

For twitches evoked by motor nerve stimulation, the time-to-peak primarily reflects processes involving dynamic properties of the muscle fibres including excitation-contraction coupling and calcium kinetics (such as release and sequestering of Ca^2+^ at the sarcoplasmic reticulum). Alternatively, twitches evoked by motor cortical stimulation reflects the conduction time along the corticospinal tract in addition to the time taken for the muscle to develop force (32–34). Our findings demonstrate that the time-to-peak twitch forces evoked by TMS, and motor nerve stimulation were comparable between groups following normal muscle contraction. However, following the exercise-induced fatiguing task, MS participants had prolonged time-to-peak in twitch force evoked by both motor cortical and motor nerve stimulation at the fatigued MVC. Previous studies in the lower limb have found that MS causes prolonged time-to-peak at rest, which were further exacerbated by exercise (35, 36). These impairments in the speed of force development of muscles of the lower limb have been suggested to be due to deconditioning or disuse (35). However, fewer impairments in muscle properties are observed for the upper body, which has been speculated as due to a lower risk of pyramidal tract damage for shorter axons supplying the upper extremity (36). Regardless of mechanism, the findings in the current study indicate that fatiguing exercise not only impairs voluntary drive from the motor cortex in people with MS, but the contractile properties of the muscle are also affected in this population.

## Conclusion

People with relapsing-remitting MS demonstrated heightened performance fatigability during sustained maximal fatiguing contractions of the elbow flexors which was associated with significantly lower biceps EMG amplitude and motor cortical voluntary activation. These findings provide evidence that MS-related performance fatigability during maximal sustained contractions is due to the inability for the motor cortex to drive the muscle, with possible contributions from altered contractile properties in the MS muscle. Deficits observed in motor cortical voluntary activation in MS were related to disease duration, and not subjective levels of fatigue (i.e. the FSS).

